# A mechanical cusp catastrophe imposes a universal developmental constraint on the shapes of tip-growing cells

**DOI:** 10.1101/2020.03.30.017095

**Authors:** Jacques Dumais, Enrique R. Rojas

## Abstract

Understanding the mechanistic basis for cell morphology is a central problem in biology. Evolution has converged on tip growth many times, yielding filamentous cells, yet tip-growing cells display a range of apical morphologies. To understand this variability, we measured the spatial profiles of cell-wall expansion for three species that spanned the phylogeny and morphology of tip-growth. Profiles were consistent with a mechanical model whereby the wall was stratified and stretched by turgor pressure during cell growth. We calculated the spatial profiles of wall mechanical properties, which could be accurately fit with an empirical two-parameter function. Combined with the mechanical model, this function yielded a “morphospace” that accounted for the shapes of diverse tip-growing species. However, natural shapes were bounded by a cusp bifurcation in the morphospace that separated thin, fast-growing cells from (nonexistent) wide, slow-growing cells. This constraint has important implications for our understanding of the evolution of tip-growing cells.

## Introduction

Cells display a vast range of morphologies that are often central to their function. For example, the filamentous morphology of tip-growing cells (Fig. 1A), which are found across the tree of life, results from the polar cell growth (Fig. 1B,C) that these cells use to explore their environment. While evolution has converged on tip growth and, *de facto*, filamentous morphology many times, the apical morphology of tip-growing cells is highly variable (Fig. 1A,S1A). It is unknown whether fitness advantages are conferred by specific apical morphologies or whether there are evolutionary or developmental constraints on the range of natural morphologies. Thus, we sought to compare the shapes of tip-growing cells observed in nature to the space of *possible* shapes, given the mechanism(s) of morphogenesis in these cells.

**Fig. 1.**
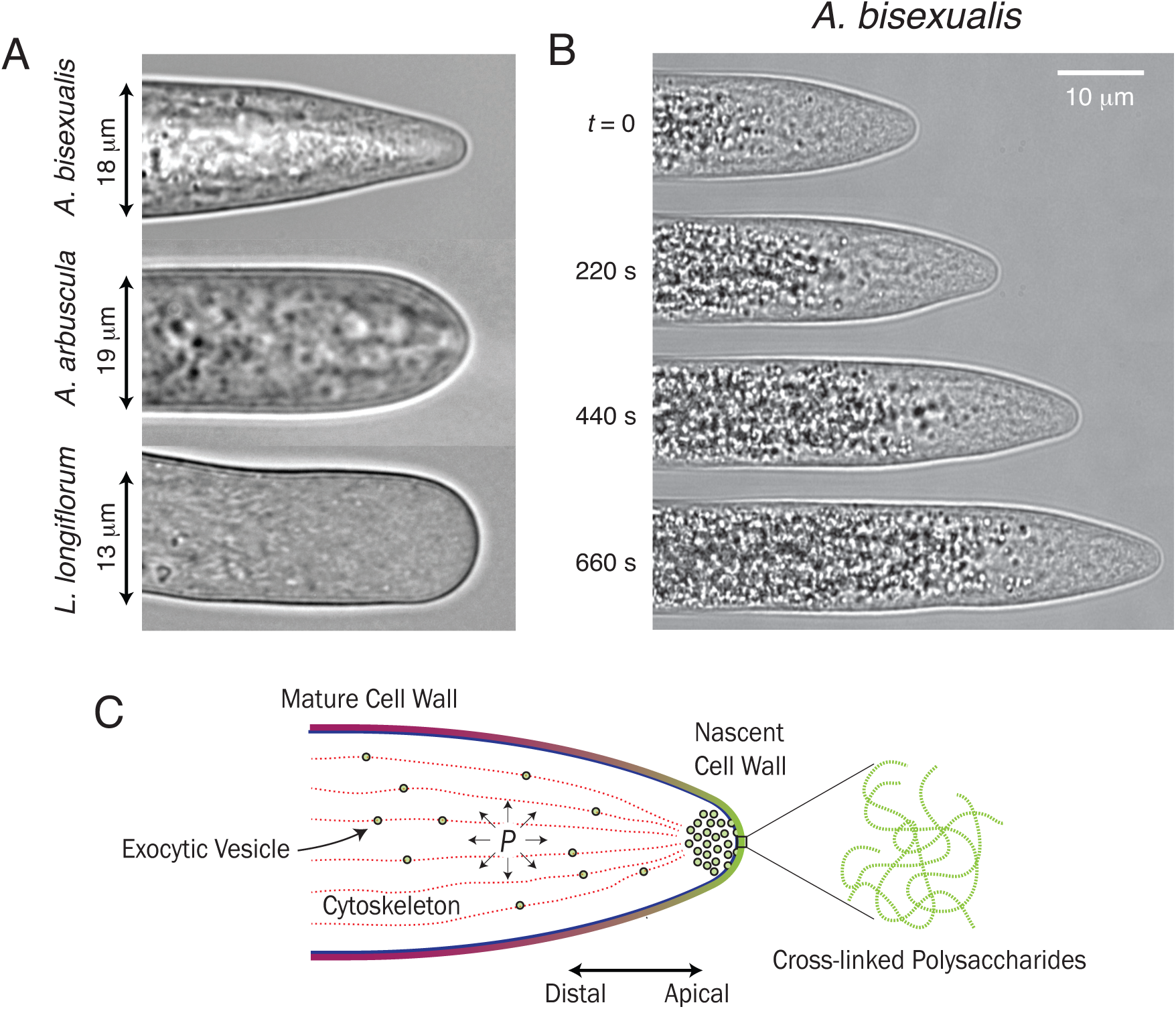
Tip growing cells are diverse in phylogeny and morphology. (A) Phase-contrast micrographs of three tip-growing cells representing protists (*A. bisexualis*), fungi (*A. arbuscula*), and plants (*L. longiflorum*). (B) Time-lapse micrograph of *Achlya bisexualis* elongating. (C) Schematic of the general features of tip-growth.

As for all walled cells, cell morphology in tip-growing cells is defined by the cell-wall geometry. Therefore, to understand cell morphogenesis it is necessary to understand the mechanism(s) by which the cell wall attains its geometry. The walls of all tip-growing cells (including those from plants, fungi, bacteria, and protists) are composed of cross-linked polysaccharides (Fig. 1C), though the specific chemistry of these polymers depends on species. Wall polysaccharides are either synthesized in the cytoplasm and then trafficked within secretory vesicles to the cell apex, where they are deposited via exocytosis into the existing cell wall (Fig. 1C), or polymerized at the plasma membrane and extruded directly into the cell wall.

In most species, the mechanism by which wall deposition is coupled to wall expansion during cell growth is unknown. However, in several tip-growing species it is clear that the biochemical composition of the cell wall is spatially graded along the apical-distal axis [1, 2], which also corresponds to the age of the cell wall since new wall material is deposited at the apex (Fig. 1C). Furthermore, in pollen tubes, a type of tip-growing plant cell, deposition of new wall material has been directly linked to enzymatic softening of the existing cell wall at the cell apex [1] and expansion of the cell wall, in turn, has been shown to depend on its irreversible mechanical deformation by the hydrostatic turgor pressure within the cell [3, 4, 5] (Fig. 1C). Similarly, in mating projections of budding yeast, cell-wall assembly and mechanical expansion are linked through a negative feedback loop that stabilizes cell growth and morphology [6].

Here, we precisely quantified the deformation of the cell wall during cell growth of two tip-growing species, a fungus and a protist (Fig. 1A). We found not only that the same constitutive mechanical model that explained cell-wall expansion in pollen tubes [5] could also do so in the non-plant species, but also that a common empirical function could account for the spatial dependence of the wall’s mechanical properties. Using these two results, we computationally described the entire range of possible cell morphologies, given the common mechanism of tip growth, and compared it to the range of tip-growing morphologies observed across the tree of life. Interestingly, natural morphologies occupied a relatively small region of the possible “morphospace.” Further computational analysis revealed that natural morphologies were bounded by an emergent cusp bifurcation, or “catastrophe,” in the morphospace that resulted from the mechanical mechanism of tip growth. We discuss the possible evolutionary reasons why this bifurcation provides such a developmental constraint.

## Results

Our goal was to determine the mechanism(s) of cellular morphogenesis across tip-growing species, determine the possible shapes that this mechanism could yield, and compare these shapes to those observed in nature. To determine the mechanism of morphogenesis, we first tested the theory of turgor-driven cell-wall expansion across a diversity of tip-growing species. To do so, we first formulated the simplest possible mechanical model of the cell wall that was consistent with our knowledge of cell-wall synthesis and deposition. Because the cell wall is invariably ≳100 nm thick, and new wall material is deposited or synthesized at the plasma membrane, the apical cell wall is stratified [7], with older strata residing toward the outer surface and younger strata toward the plasma membrane. This precludes a fully isotropic cell wall (Fig. 2A), but is consistent with a “transversely isotropic” cell wall, where within a given stratum, wall material has no preferred orientation parallel to the plane of the wall (Fig. 2A). Mathematically, this symmetry within the wall corresponds to a simple constitutive relationship between the principal in-plane wall expansion rates,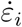, and tensions, λ_*i*_ (Fig. 2B):

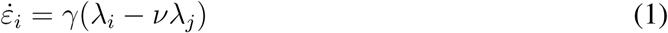

where *v* is the “flow coupling” and γ = (*ηδ*)^*-*1^ is the “surface extensibility,” equal to the reciprocal of the product of wall viscosity, η, and wall thickness, δ. The indices *i, j* refer to orthogonal principal directions within the plane of the cell wall. The quantities ε_*i*_, λ_*i*_, and γ, are functions of space on the apical cell surface, which can be parameterized by the arclength, s, away from the cell pole (Fig. 2A).

**Fig. 2.**
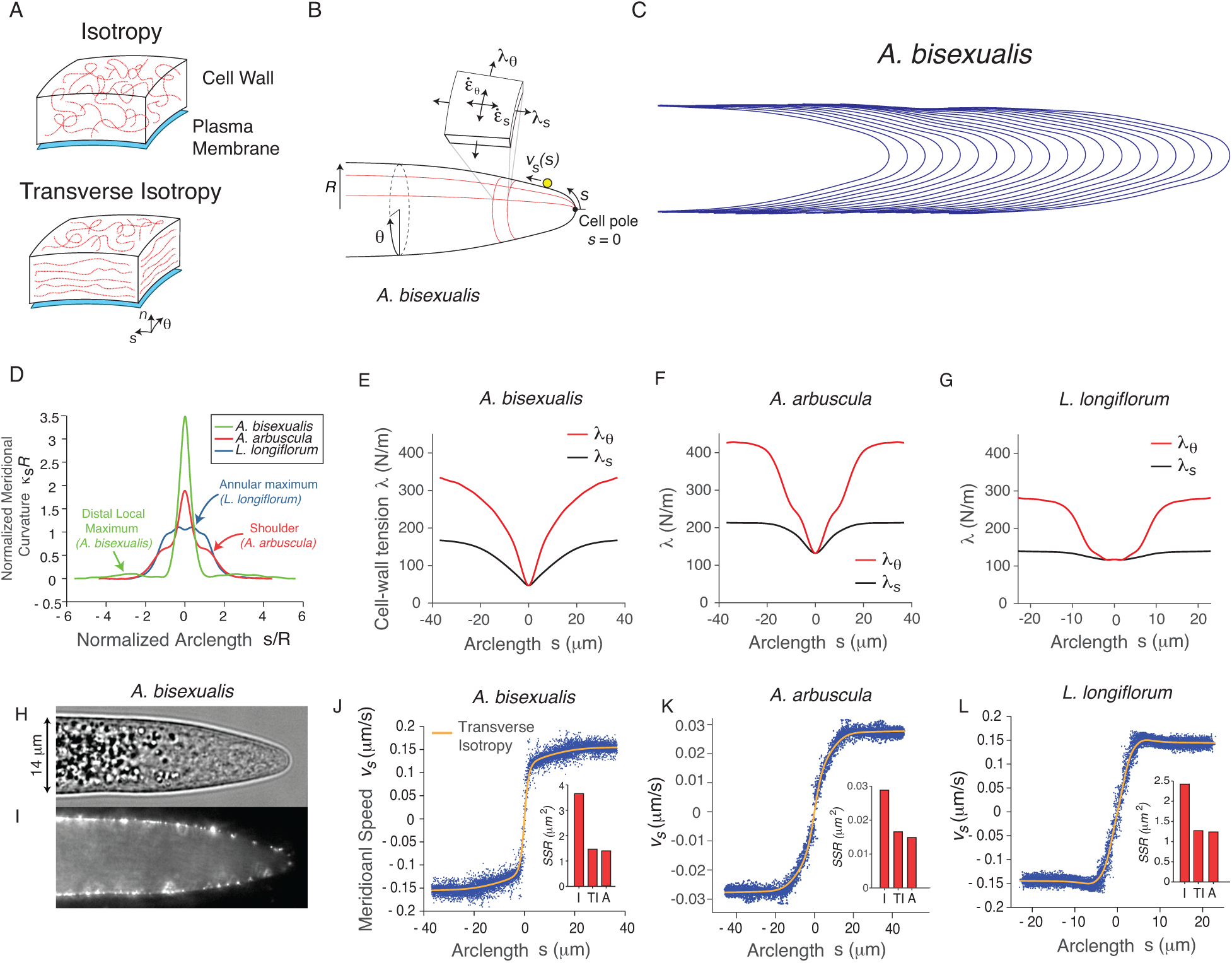
A common mechanical mechanism drives cell-wall expansion in diverse tip-growing species. (A) Illustration of symmetries within a cell wall with isotropic and transversely isotropic material properties. Red dotted lines represent cell wall polymers. (B) Parameterization of the cell surface and illustration of key variables. (C) Computationally extracted experimental cell-wall geometry from a growing *A. bisexualis* cell. (D) Representative mean meridional curvature (normalized by cell radius) versus normalized arclength for cells from three species. (E-G) Representative spatial profiles of principal cell-wall tension versus arclength for *A. bisexualis* (*n* = 2 cells), *A. arbuscula* (*n* = 4 cells), and *L. longiflorum* (*n* = 4 cells). (H) Phase-contrast image of *Achlya bisexualis*. (I) Epifluorescence micrograph of the same cell in (H) showing fluorescent microspheres attached to the cell wall. (J-L) Meridional speed versus arclength for the same cells for which cell-wall tensions were calculated (E-G). The orange lines are the best fits from the transverse isotropy model. (insets) Sum of squared residuals (*SSR*) of the fits of three symmetries (isotropy (I), transverse isotropy (TI), anisotropy (A)) tested against the meridional speed profiles.

Transverse isotropic properties explain the spatial dependence of cell-wall expansion in root hairs [4] and pollen tubes [5]. To test whether this mechanical model could account for cell-wall expansion dynamics across a range of tip-growing cells, particularly non-plant species, we measured the principal wall tensions and expansion rates in the hyphae of *Achlya bisexualis* (a protist, Fig. 1A) and *Allomyces arbuscula* (a fungus, Fig. 1A); together with lily pollen tubes (Fig. 1A) these systems spanned tip-growing phylogeny and morphology (Fig. 1A, S1A). To measure the principal wall tensions, we recorded time-lapse micrographs of cells as they grew (Movie S1), computationally tracked apical morphology from these micrographs (Fig. 2C) and then calculated the average meridional curvature of the cell wall (Fig. 2D), from which the spatial dependence of the tensions (Fig. 2E-G) could be calculated via force balance equations (*Methods*, [4]).

To measure the principal expansion rates, we adhered electrically charged, fluorescent microspheres to the cell walls (Fig. 2H,I) and recorded time-lapse micrographs of them as the cells grew (Movie S2). In the frame of reference of the cell pole, these microspheres move along cell meridians with speed *v*_*s*_(*s*) (Fig. 2J-L,S2). Along with cell-wall geometry (Fig. 2D,S1B), the profile of *v*_*s*_(*s*) completely describes cell-wall expansion. That is, the principal expansion rates, 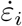, can be calculated directly from these variables [8].

We found that for each species, the transverse isotropic mechanical model (Eq. 1) provided an excellent fit of *v*_*s*_(*s*) (Fig. 2J-L). Conversely, if the wall was assumed to be fully isotropic, the model provided a poor fit, while increasing the model’s fitting power by relaxing the constraint of transverse isotropy (i.e., assuming full anisotropy) only slightly improved its fit (Fig. 2J-L, inset), suggesting that a common mechanical mechanism, that of transversely isotropic mechanical expansion of the cell wall, underlies cellular morphogenesis across a wide diversity of tip-growing species. This precise analysis is supported by several previous low-resolution measurements of cell-wall expansion in diverse tip-growing systems that show that the circumferential expansion rate is higher than the meridional expansion rate across the entire apical cell wall except for near the pole [4, 5, 9, 10]; this feature results from mechanical cell-wall expansion since the principal cell-wall tensions have the same anisotropy due to the generic tip-growing cell morphology [11].

This common mechanical model gives rise to specific morphologies via the “surface extensibility profile,” *γ*(*s*) (Eq. 1). Thus, to examine the space of possible shapes yielded by the model, we first calculated this variable for each of the three species by inputting our experimental tension and expansion-rate profiles (Fig. 2E-G,J-L) into Eq. 1 (*Methods*). The experimental surface extensibility profiles qualitatively resembled the curvature profiles for each species, respectively: the *A. bisexualis* and *A. arbuscula* profiles displayed sharp central peaks while the *L. longiflorum* profile had a slight annular maximum (Fig. 3A-C). Both the *A. bisexualis* and *A. allomyces* profiles had apparent “long tails.” As such, the profiles of each species could be accurately fit with an empirical function that featured two length scales, in particular a superposition of two Gaussian functions: 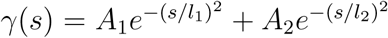 (Fig. 3D). Computationally simulating cell growth using the transverse isotropy model combined with the empirical “Gaussian mixture” surface extensibility profile accurately predicted apical cell morphology (Fig. 3E-G, Fig. S3A-D). Furthermore, cells from a given species clustered with respect to the the parameters of the Gaussian mixture model that best fit their surface extensibility profile (Fig. S3E).

**Fig. 3.**
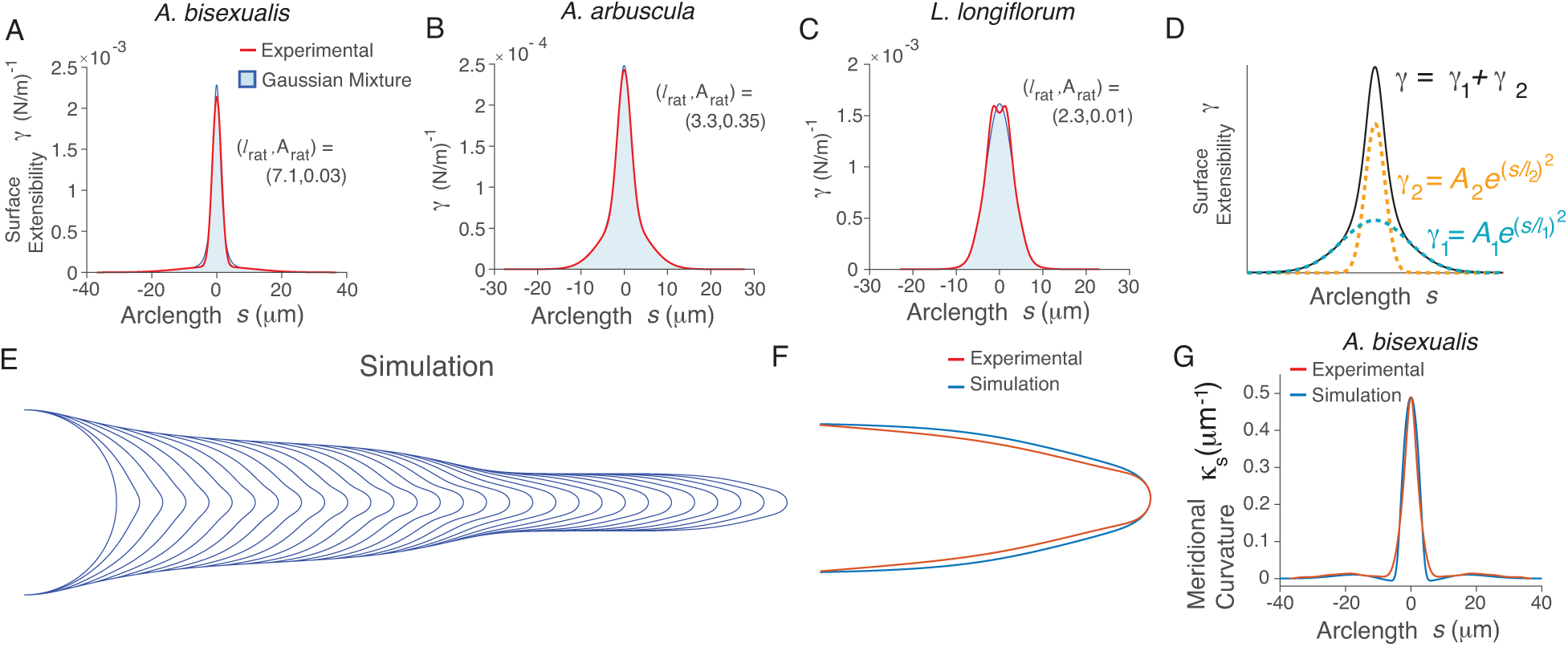
A two-parameter surface extensibility profile can account for morphological diversity in tip-growing cells. (A-C) Experimentally determined surface extensibility profiles for cells for which meridional speed profiles were measured (red lines). The light blue areas show the best fit by the empirical “Gaussian mixture” function. The values of (*l*_rat_, *A*_rat_) that yielded the best fit are given. (D) Illustration of the empirical “Gaussian mixture” function used to fit the extensibility profiles. (E) Computational simulation of cell growth using the transverse isotropy model and an empirical Gaussian mixture surface extensibility profile with parameters that yielded the best fit of the experimental surface extensibility profile of the *A. bisexualis* cell shown in (A). The simulation was initiated with a hemispherical geometry. (F) Comparison of the apical geometry of of the simulation shown in (E) to the experimental apical geometry for the *A. bisexualis* cell that was being simulated. (G) Comparison of the meridional curvature profile of the simulated cell shown in (E) to experimental profile of the *A. bisexualis* cell that was being simulated.

Given this result, we asked whether the diversity of tip-growing morphologies observed across the tree of life could be fit by the transverse isotropy mechanical model with a Gaussian mixture surface extensibility profile. To answer this, we first quantified the apical morphologies of a wider diversity of tip-growing cells from various taxa (Fig. S1). To objectively quantify the range of apical morphologies, we used principal components analysis (PCA, Fig. 4A-D), which revealed that the first principal variable, P1, could account for ≈ 95% of shape variation across this data set (Fig. 4C), demonstrating that this scalar is a useful proxy for cell shape. We then performed a parameter-space search across the three parameters in the model that dictated apical morphology. While the model has five free parameters, cell size and cell elongation rate can be fit independently of apical morphology, leaving three free parameters: *l*_rat_ = *l*_1_/*l*_2_, *A*_rat_ = *A*_1_/*A*_2_, and *v* (*Methods*). Most of the “morphospace” spanned by *l*_rat_ and *A*_rat_ had high values of P1, yielding relatively oblate pollen tube-like apical geometries (Fig. 4E,F). However, there was a narrow region of morphospace that displayed lower values of P1, corresponding to the more tapered “*Achlya*-like” apical geometries (Fig. 4E,F). Variations in the flow coupling, *v*, affected the position of the region of low P1 values, but not the qualitative dependence of P1 on *l*_rat_ and *A*_rat_ (Fig. S3F-K). Away from the region of low P1 values, for a given surface extensibility profile, flow coupling slightly affected the cell radius but had little effect on cell morphology (Fig. S3J).

**Fig. 4.**
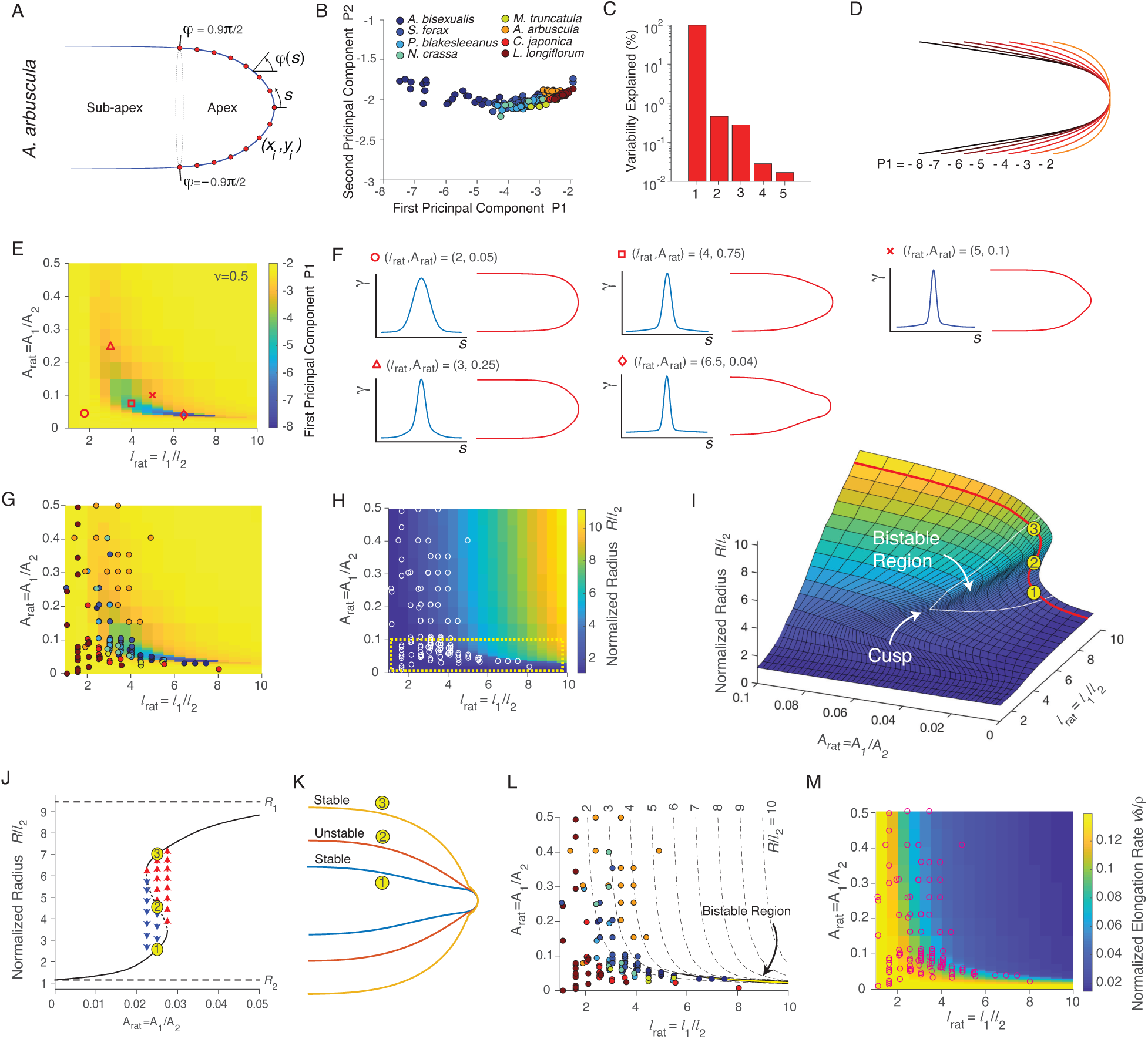
A shape instability constrains observed cell shapes. (A) Time-averaged cell outline extracted from an *A. arbuscula* cell, including a subset of the discrete co-ordinates used for PCA analysis (red dots). The apical cell wall was defined as the portion corresponding to *-*0.9*π*/2 < *φ*(*s*) < 0.9*π*/2, where *φ* is the angle between a vector normal to the cell surface and the cell axis. (B) Experimental values of the first two principal components *P*1 and *P*2 for each cell from a diversity of tip-growing organisms (*n* = 32, 23, 24, 4, 7, 36, 32, 25 for *A*.*b*., *S*.*f*., *P*.*b*., *N*.*c*., *M*.*t*., *A*.*a*., *C*.*j*., *L*.*l*., respectively). (C) Percent of morphological variation explained by the first five principal components. (D) Apical geometry versus *P*1. For this calculation, the remaining principal variables were held constant, with each one equal to the average experimental value across all cells from all species. (E) P1 of simulated cells using the empirical Gaussian mixture surface extensibility profile (Fig. 3D), as a function of *l*_rat_ and *A*_rat_. A flow coupling of *v* = 0.5 was used. (F) Surface extensibility profiles and apical geometries from five selected (*l*_rat_, *A*_rat_) co-ordinates within the morphospace, indicated by red shapes in (E). (G) Values of (*l*_rat_, *A*_rat_) that, when used to simulate cell growth, yielded the best fit of the geometry for each cell of each species, overlaid on the values of *P*1. The color code for species is the same as that used in Fig. 4B. (H) Radius of simulated cells, normalized by *l*_2_, as a function of *l*_rat_ and *A*_rat_. The white circles indicate the co-ordinates that yielded the best fit of the experimental morphologies shown in (G). (I) Radius of simulated cells, normalized by *l*_2_, as a function of *l*_rat_ and *A*_rat_, within the region enclosed in the yellow dotted box in (H). (J) Normalized radius as a function of *A*_rat_ for *l*_rat_ = 9.5. This slice is shown by the red line in (I). The dotted section of the curve represents unstable solutions. The dashed lines indicate the radii *R*_1_ and *R*_2_ of simulated cells with a single Gaussian profile of *l* =1 and *l* = 9.5, respectively. The arrows indicate the convergence of the radius to that of the stable shape solution when a simulation of cell growth was initiated with shape that was was a linear superposition of the stable and unstable solution with radius given by the co-ordinate of the arrow. (K) Steady-state geometries of cells with (*l*_rat_, *A*_rat_)= (9.5, 0.025), including 2 stable geometries and 1 unstable geometry. The co-ordinates of the solutions are indicated by the yellow points in (I) and (J). (L) Values of (*l*_rat_, *A*_rat_) that, when used to simulate cell growth, yielded the best fits of the geometry for each cell of each species, overlaid onto contours of the normalized radius of simulated cells as a function of *l*_rat_ and *A*_rat_. (M) Elongation rate, normalized by *δ*/*ρ*, as a function of *l*_rat_ and *A*_rat_. The white circles indicate the co-ordinates that yielded the best fit of the experimental morphologies shown in (G) and (L).

We used a least-squared fit method to find the parameters in the morphospace that yielded the best fit of each cell from each tip-growing species. The morphospace provided a highly accurate fit of the apical morphology in all cases (Fig. S3A-D,L,M, *Methods*). Interestingly, we found that the values of *l*_rat_ and *A*_rat_ that provided the best fit across species occupied a limited region of the morphospace, corresponding to low values of at least one of these variables, and appeared to be bounded by the narrow region of low P1 (Fig. 4G).

In order to understand why the region of low P1 values provided such a strong constraint on the observable shapes of tip-growing cells we analyzed other properties of cell geometry as a function of *l*_rat_ and *A*_rat_. We discovered that for a given value of *l*_2_ (the length scale of the narrower peak in the surface extensibility profile; Fig. 3D), that cell radius varied as a function of *l*_rat_ and *A*_rat_, with lower values of these variables yielding thinner cells (Fig. 4H). Given that low values of *l*_rat_ and *A*_rat_ also provided the best fits to empirical cell shapes (Fig. 4G), this implies that cells from across nature are relatively thin compared to the length scale of their surface extensibility profile. While the transition between thin cells to wide ones was gradual as *l*_rat_ was increased, it was extremely sharp as *A*_rat_ was increased for *l*_rat_ ≳ 5 (Fig. 4H). A closer computational examination of this region of the morphospace revealed that this sharp boundary corresponded to the bistable region of a cusp bifurcation or “catastrophe” [12] (Fig. 4I). That is, above a critical value of *l*_rat_ the surface of stable solutions folds such that there is a narrow region of the morphospace where there are two stable solutions (and one unstable solution) for cell shape for a single surface extensibility profile (Fig. 4I,J). For computational simulations of cell growth in this region of the morphospace, the steady-state shape depended on the shape from which the simulation was initiated (Fig. 4J,K). With respect to cell shape, one branch of the bistable solution corresponded to tapered, *Achlya*-like cell geometries, whereas the other branch corresponded to more oblate cells, but with a small region of high curvature near the cell pole (Fig. 4J,K). Remarkably, whereas natural cell shapes were generally confined to low-radius regions of the morphospace, tapered cell shapes were strictly confined to the low-radius side of the bistable region (Fig. 4L). Thus, the widespread mechanical mechanism of tip growth results in a shape instability and, empirically, we find that this instability imposes a universal developmental constraint on apical shapes of tip-growing cells. We discuss the implications of this constraint below.

## Discussion

We discovered an emergent shape instability in tip-growing cells that results from the mechanical mechanism of tip growth and that imposes a strong developmental constraint on the shapes of these cells from across nature. Given this constraint, there is some flexibility with respect to shapes that cells can assume, however, there appears to be a trade-off between flexibility in shape and flexibility in radius (relative to the length scale of the surface extensibility profile, *l*_2_). That is, away from the instability, while cells are still confined to the low-radius region of the morphospace, radius changes gradually with *l*_rat_ (Fig. 4H) and therefore there is some flexibility in terms of the range of radii cells can assume, with limited flexibility in terms of shape (e.g., *A. arbuscula*, Fig. 4G). On the other hand, species that exhibit a wider range of shapes (from oblate to tapered) must assume their tapered shapes near the instability, which provides a strong constraint in terms of cell radius (e.g. *A. bisexualis*, Fig. 4G,H).

While the cusp catastrophe provides a clear developmental constraint, it is unclear why this is so. The elongation rate of the cell scales inversely as the radius (*v* = *ρ*/*Rδ*, where *ρ* is the total rate of cell wall biosynthesis and *δ* is the wall thickness), and therefore low values of *l*_rat_ and *A*_rat_ yield faster-growing cells for a given rate of material addition (Fig. 4M). Thus, it is reasonable to view the dependence of elongation rate on *l*_rat_ and *A*_rat_ as a fitness landscape, whereby fast-growing cells outcompete slower-growing ones. Tip growth is a mechanism by which cells can “move” in lieu of true cellular motility: for the hyphae of fungi and water molds, as well as root hairs, this form of growth serves to explore the environment in search of nutrients, while for pollen tubes the object is to fertilize the ovule. In each of these cases, fast elongation (and therefore thin cells) will be selected by nature. Minimizing the ratio between surface area and volume by maintaining thin cells may also be important for nutrient uptake. Since the range of shapes permitted by the mechanical mechanism of morphogenesis is correlated with the relative growth rate (Fig. 4M) and surface-area-to-volume ratio (via cell radius, Fig. 4H), nature would favor certain shapes.

However, for several reasons it is difficult to explicitly test that the cusp catastrophe represents a fitness landscape. The primary reason is that we expect there to be low variability in relative cell radius (*R/l*_2_) and relative elongation rate (*v* = *ρ*/*Rδ*) for the very reason that tip-growing species have evolved to optimize these variables. Since there is, however, a limited range of elongation-rate variation across and within tip-growing species, it would be useful to compare this to cell shape. However, comparing the elongation rate between species is not useful because the species we assayed inhabit a wide range of environmental niches, and many other ecological and evolutionary factors may influence their absolute growth rate. Within a species, it would be useful to examine trade-offs between shape and elongation rate, but from our model, we can only calculate the elongation rate relative to the cell-wall biosynthesis rate, *ρ*, which is not currently possible to measure. This rate, as well as the length scale of cell-wall surface extensibility could depend on cell radius [13], which would confound this analysis. Future research should address these measurements.

Though the cusp catastrophe provides a proximate developmental constraint, it emerges from the underlying transverse isotropical expansion of the cell wall. Why is this material symmetry within the cell wall so widespread in nature? Transverse isotropy *itself* may be a developmental constraint. This symmetry follows from the cell-wall stratification that, in turn, results from continuous exocytosis of new amorphous wall polymers at the inner face of the cell wall and subsequent stretching of these layers. In order to violate transverse isotropy, the cell would either need to i) “mix” strata within the cell wall after exocytosis, so as to obtain an isotropic cell wall, or ii) deposit highly ordered wall polymers within strata so as to obtain a fully “plane anisotropic” cell wall. Cell-wall mixing seems impossible, and exocytosis of amorphous material cannot confer directional order. On the other hand, ordered deposition of wall polymers occurs in diffusely growing cells but has not been observed in tip-growing cells. Ordered deposition requires that the complexes that perform this deposition be able to sense their location and orientation within the cell (via, for example, curvature sensing of the cell surface), which may be prohibitively difficult within the apical geometry of a tip-growing cell compared to the regular cylindrical geometry of a diffusely growing cell.

While we included a diversity of tip-growing cells in our study, there is a conspicuous omission: the bacteria. It was commonly assumed that bacteria also use turgor to drive cell-wall expansion, which could have provided a unifying evolutionary history for this mechanism in other kingdoms. However, recent studies suggested that turgor is only important for wall expansion in Gram-positive organisms [14, 15], which comprise only two, relatively small clades compared to Gram-negative organisms [16]. Like higher walled organisms, Gram-positive bacteria possess a relatively thick (≳20 nm) cell wall [17], whereas Gram-negative organisms possess a thin cell wall, perhaps as thin as a molecular monolayer [18]. That is, the mechanism for turgor-driven growth may have co-evolved with cell-wall thickness multiple times. Due to their small size, the actinomycetes, a phylum of tip-growing Gram-positive bacteria, have traditionally been overlooked in experimental biomechanics studies. However, measurement of the wall expansion rate in rod-shaped bacteria has now been achieved [19], and similar methods may soon permit the testing of the mechanical basis of bacterial tip growth in the quest for a unified model of tip-growth morphogenesis.

## Materials and Methods

### Strains and Media

*C. japonica* was collected, stored and cultured in the same manner as *L. longiflorum*, described previously [5]. *A. arbuscula, A. bisexualis, S. ferax, P. blakeseanus* were obtained from Carolina, Inc. *N. crassa* was a gift from Anne Pringle. Fungal and oomycete cultures were propagated at room temperature on solid yeast-malt growth medium consisting of 0.3% (m/v) yeast extract, 0.3% malt extract, 0.5% peptone, 1% dextrose, and 1% agarose.

### Imaging and image analysis

Imaging of *C. japonica* was performed as for *L. longiflorum* [5]. Briefly, 1 hour prior to imaging, freeze-dried pollen grains were immersed in liquid growth medium. A thin layer (<1 mm) of solid growth medium (1% low melting point agarose) was deposited onto the bottom of a custom-made chamber, consisting of a dental polymer gasket affixed to a cover glass. While the agarose was still in a molten state, pollen grains were partially embedded in the gel by rinsing it with a suspension of grains in liquid growth medium. The slide chambers were then filled with liquid medium. Cells that emerged from grains affixed to the agarose in this way were likely to grow along the gel-liquid interface, which was ideal for imaging.

For imaging fungal and oomycete hyphae (except for *A. arbuscula*) dental-polymer slide chambers were prepared, as above, with 1% low melting point agarose yeast-malt growth medium. After the agarose solidified, the chamber was inoculated with a small block of agarose from the growing front of the mycelium in the propagation culture. The top of the slide chamber was then sealed with a cover slip. After incubation for 48 − 72 hours at room temperature, the top of the chamber was opened, and filed with liquid medium for imaging.

For imaging *A. arbuscula*, cultures were propagated in special cover-glass bottomed Petri dishes (MakTek). The bottom of the dishes were covered in a layer (≈ 2mm) of solid growth medium (1% agarose) and one edge of this substrate was inoculated. The hyphae grew in the thin layer of liquid between the cover glass and the agarose at the bottom of the dish. Cells were imaged 5 − 6 days after inoculation.

Cell tracking and calculation of principal tensions was performed using a custom MATLAB algorithm, as described previously [5].

### Principal components analysis

To perform principal components analysis, we first reconstructed the time-averaged apical geometry for each cell by integrating the time-averaged meridional curvature profile:

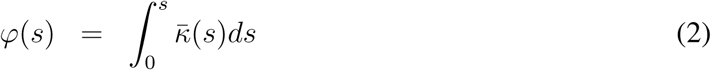

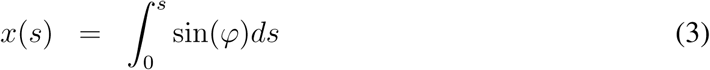

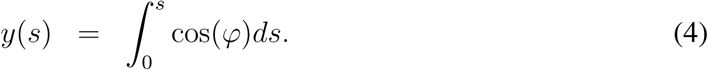

So as to only perform PCA on the apical region and not the subapical region, we truncated the outlines using *φ* = [−0.9*π*/2, 0.9*π*/2] as lower and upper bounds, and then interpolated the outlines with 75 equally spaced fiducial points. We found that *φ* = 0.9*π*/2 was the largest cutoff we could use while not including large portions of the subapical, nearly cylindrical region of the cell. The axis of the apex was found by averaging opposing fiducial points, and the cell apices were rotated such that their axes were aligned. Principal components analysis was then applied to the interpolated (*x, y*) outlines at these points using the MATLAB command *pca*.

### Microsphere imaging and measurement of meridional speed

For microsphere experiments, 0.1 *µ*m aminated fluorescent microspheres (Sigma-Aldrich) were included in all media (10^−4^ dilution from stock), including solid media. For *A. bisexualis*, just after the slide chamber was filled with microsphere-containing media prior to imaging, the microspheres had a high affinity for the cell walls. As the charges on the microspheres were neutralized, they lost their affinity for the cell wall, making it necessary to re-immerse in freshly mixed microsphere growth medium every 10 − 15 minutes during imaging.

For *A. arbuscula*, a section of the solid media several millimeters in front of the leading edge of the mycelium was excised with a scalpel, forming a well to which microsphere-containing liquid media was added. Over the course of minutes, the microspheres diffused through the liquid layer between the cover glass as the solid media and reached the mycelium, whereupon they adhered to the hyphal cell walls.

All cells were imaged through a 60× water immersion lens and a 1.6× projection lens, for a total magnification of 96×, on an inverted microscope (Olympus IX71, Center Valley, PA). For epifluorescence microscopy, the samples were illuminated with a 120 W mercury arc lamp (EXFO, Quebec, QC) and viewed through a Texas red filter. Time-lapse images were recorded with a 14-bit cooled CCD camera (The Cooke Corporation, Romulus, MI). IPLab (Scanalytics, Rockville, MD) was used for image acquisition. Acquisition frame rate was between 3 and 20 s, depending on species.

Fluorescent microsphere tracking and calculation of meridional speed was performed as described previously [5].

The fitting of the meridional speed profiles was done as described previously. For the isotropy and transverse isotropy models, the shape of the cell was fit perfectly, which predicts the meridional speed profile given a value of *v*, the flow coupling [20]. Isotropy requires that *v* = 0.5 whereas for transverse isotropy *v* is a fitting parameter. For the anisotropic model, we performed a “free fit” [4].

### Simulations of cell growth and parameter space search

Simulations of cell growth for transverse isotropy with an arbitrary extensibility profile were performed with a custom MATLAB routine. An initial hemispherical geometry was defined with *N* (an odd number typically between 101 and 301) discrete (*x, y*) co-ordinates. The arclength, s, was calculated on the geometry, with the polar point defined as s = 0 (Fig. 2B). The surface extensibility profile to be simulated, *γ*(*s*), was then prescribed on these points. The radius of the hemisphere was arbitrary but was generally chosen to be somewhat larger than the largest length scale associated with the extensibility profile.

At each step of the simulation, first the principal curvatures in the cell wall were calculated [8]:

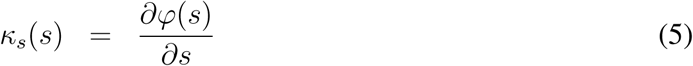

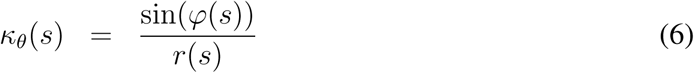

where *r*(*s*) is the radial distance from the cell axis to the fiducial point. From the principal curvatures, the principal stresses were calculated using using the force-balance equations [8]:

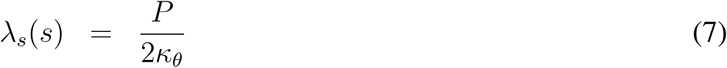

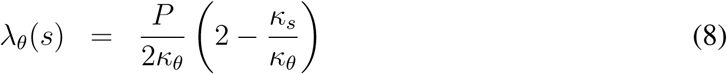

From the principal tensions, the principal expansion rates were calculated using Equation 1 of the main text:

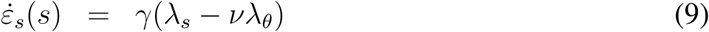

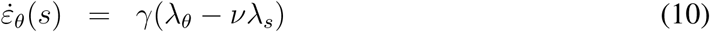

To then calculate the velocity of each fiducial point we inverted the kinematic equations [8]:

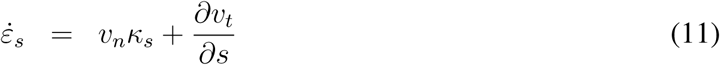

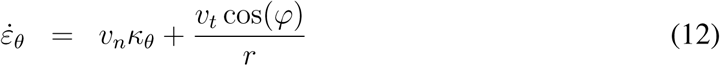

where *v*_*n*_ and *v*_*t*_ are the velocity components of the discrete points in the directions normal and tangential to the cell outline. Finally, the displacement of each point *v*_*n*_*dt, v*_*t*_*dt* was calculated (where *dt* is the time between steps), converted to cartesian co-ordinates, and added to the cell geometry to find the updated cell geometry. This updated geometry was smoothed with a smoothing spline (the MATLAB csaps command, with smoothing factor 0.9999), which was critical to prevent small errors from causing instabilities. This routine was iterated until cell geometry converged. At each step, the time step was updated to ensure that the displacement at the pole converged to a fixed, small value.

To validate the simulation routine, we input the experimental surface extensibility profiles, which were calculated to perfectly fit the experimental cell geometry given the transverse isotropy model. In our simulations, this surface extensibility profile yielded a meridional curvature profile that was indistinguishable from the experimental profile (data not shown).

Similarly, we explicitly showed that while scaling of the height or width of a given surface extensibility profile affected the size and elongation rate of the simulated cells, it had no effect on cell shape. When cell size was normalized for a surface extensibility profile that had been re-scaled, it showed an indistinguishable meridional curvature profile to a cell simulated using the raw surface extensibility profile (data not shown).

For the parameter space searches described in Fig. 4E and Fig. S3I-L, *l*_rat_ was varied on the domain (1, 10) at 0.25 increments and *A*_rat_ was varied on the domain (0, 0.1) at 0.005 increments and the domain (0.1, 0.5) at 0.05 increments. For each value of *A*_rat_, the search was initiated at *l*_rat_ =1 and *A*_rat_ was increased incrementally to 0.5, using the solution of cell shape from the previous parameter co-ordinate as the initial condition for each simulation. The fold bifurcation was not evident in these parameter space searches because we only increased *A*_rat_ and did not decrease it.

To resolve the fold bifurcation shown in Fig. 3I, we focused on the domain *l*_rat_ ∈ (1, 10) (increments of 0.25) and *A*_rat_ ∈ (0, 0.1) (increments of 0.0025). We searched the space as before, but both increased and decreased *A*_rat_ for a given *l*_rat_. To resolve the fold bifurcation we found (*l*_rat_, *A*_rat_) co-ordinates where there were two stable solutions. For each one of these co-ordinates, from the two stable apical geometries,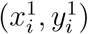 and 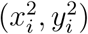, we found ten intermediate geometries 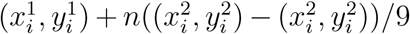, where *n* = 0, 1, …, 9. We then initiated a simulation with each of these geometries and determined to which stable solution the simulation converged (Fig. 4J). The unstable solution (Fig. 4K) was computed to be the mean of neighboring geometries that converged to different stable fixed geometries (Fig. 4J). From these unstable geometries, the radius could be calculated to find the whole manifold of solutions shown in Fig. 4I.

To calculate the best fit of the model for each experimental cell geometry, we first found the time-averaged cell outline, which was found by integrating the time-averaged meridional curvature profile using Equations 2-4. We discretized the time-averaged experimental cell outline into the same number of points that was used to simulate tip growth in the computational parameter space search. We then calculated the normalized average error,

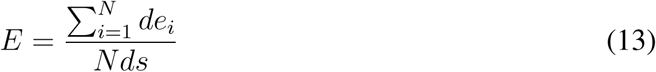

which is the average error of each discretized point relative to the interpoint arclength. We found values of *l*_rat_ and *A*_rat_ that minimized this error.

## Acknowledgments

This work was funded by the Human Frontier Science Program (RGP0018/2006-C) and the NYU Department of Biology.

## Supplementary Information

### Supplementary Figure Legends

**Fig. S1.**
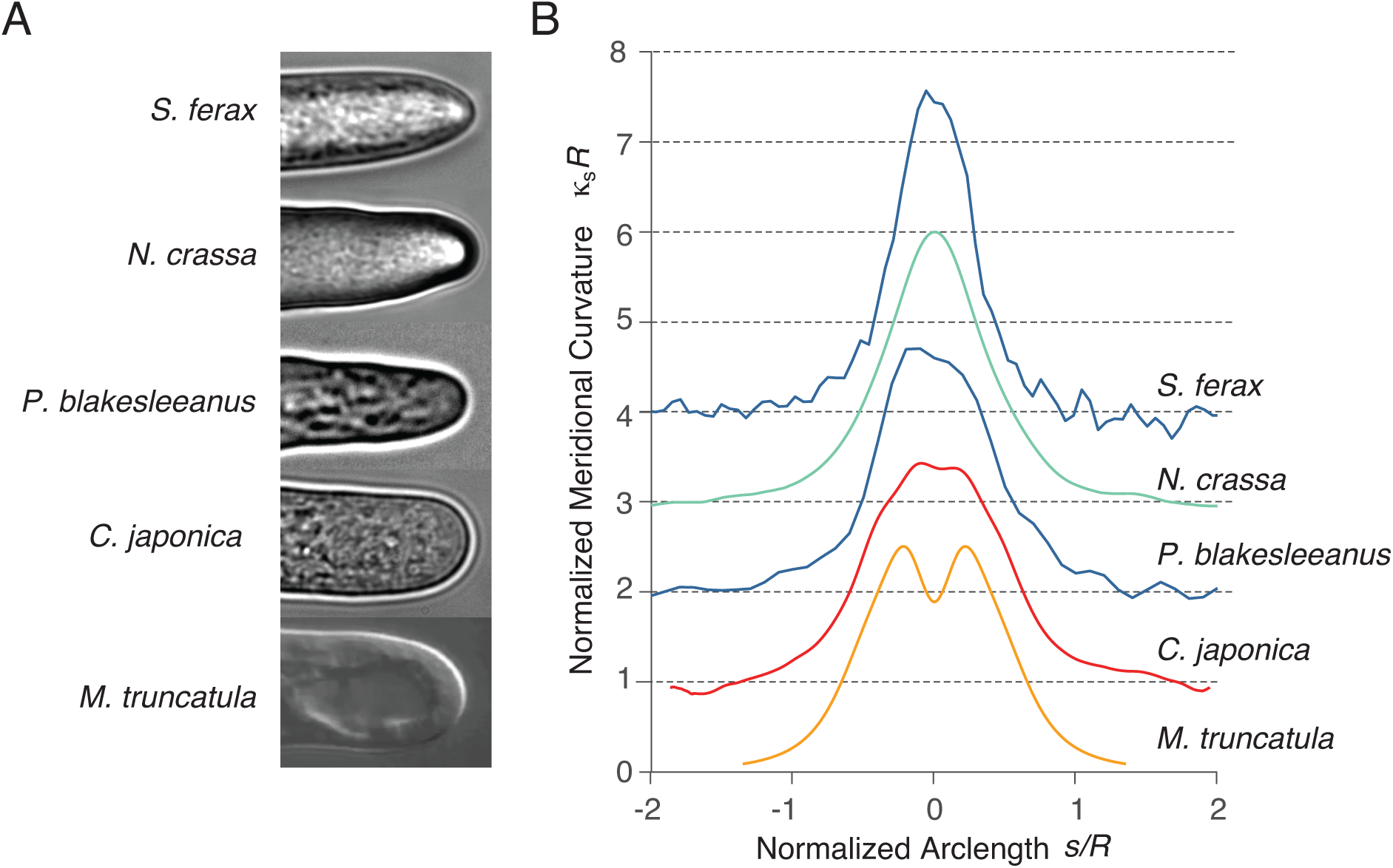
Apical geometry varies across the tree of life. (A) Phase micrographs of five tip-growing organisms. (B) Representative mean meridional curvature (normalized by cell radius) versus normalized arclength for cells from the species shown in (A).

**Fig. S2.**
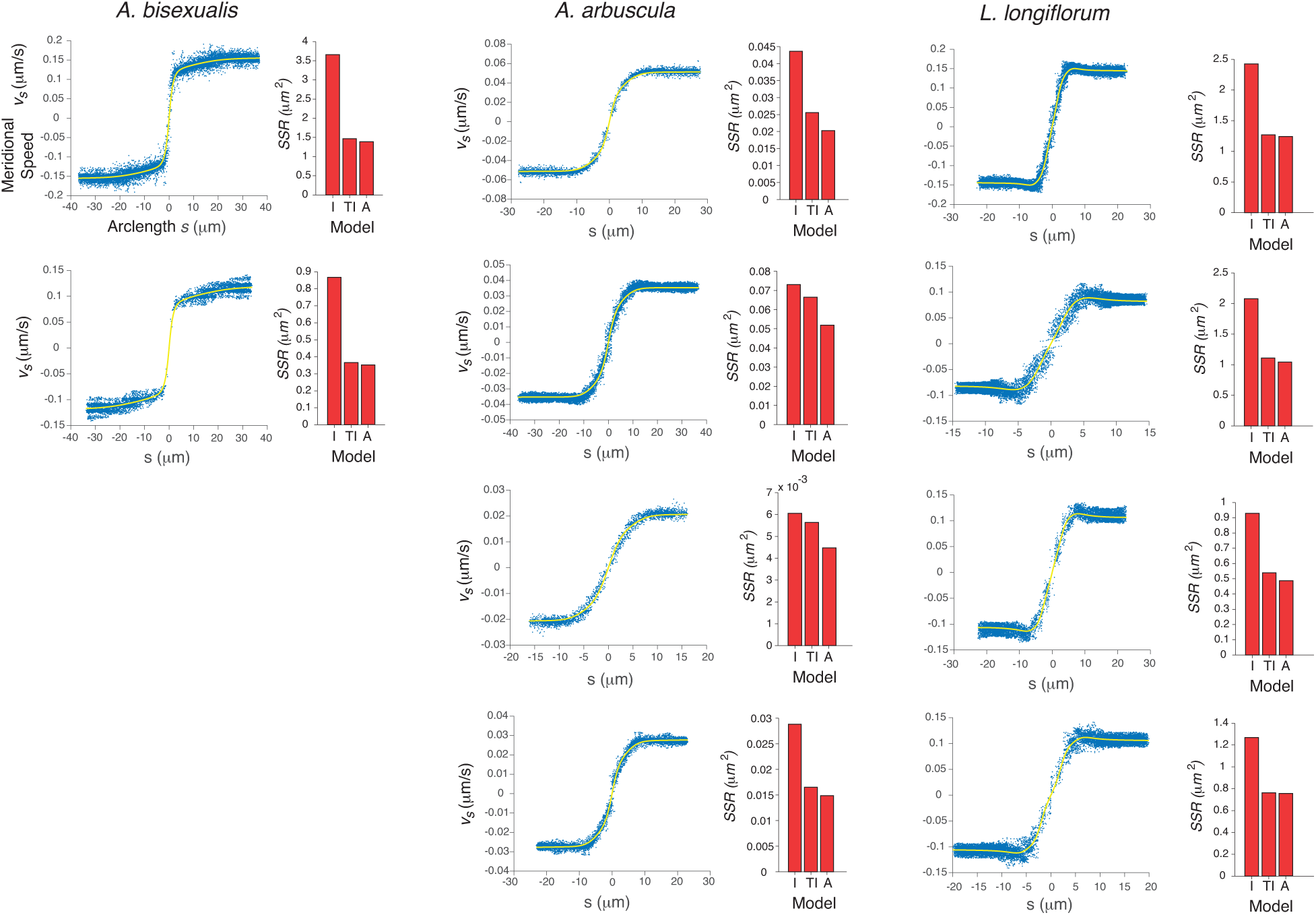
Transverse isotropy predicts the mechanics of cell-wall expansion. Meridional speed versus arclength for *A. bisexualis* (*n*=2 cells), *A. arbuscula* (*n*=4 cells), and *L. longiflorum* (*n*=4 cells). The yellow lines are the best fit with the transverse isotropy model. Next to each profile is the sum of squared residuals (SSR) for the best fits from the three mechanical models: isotropy (I), transverse isotropy (TI), and anisotropy (A).

**Fig. S3.**
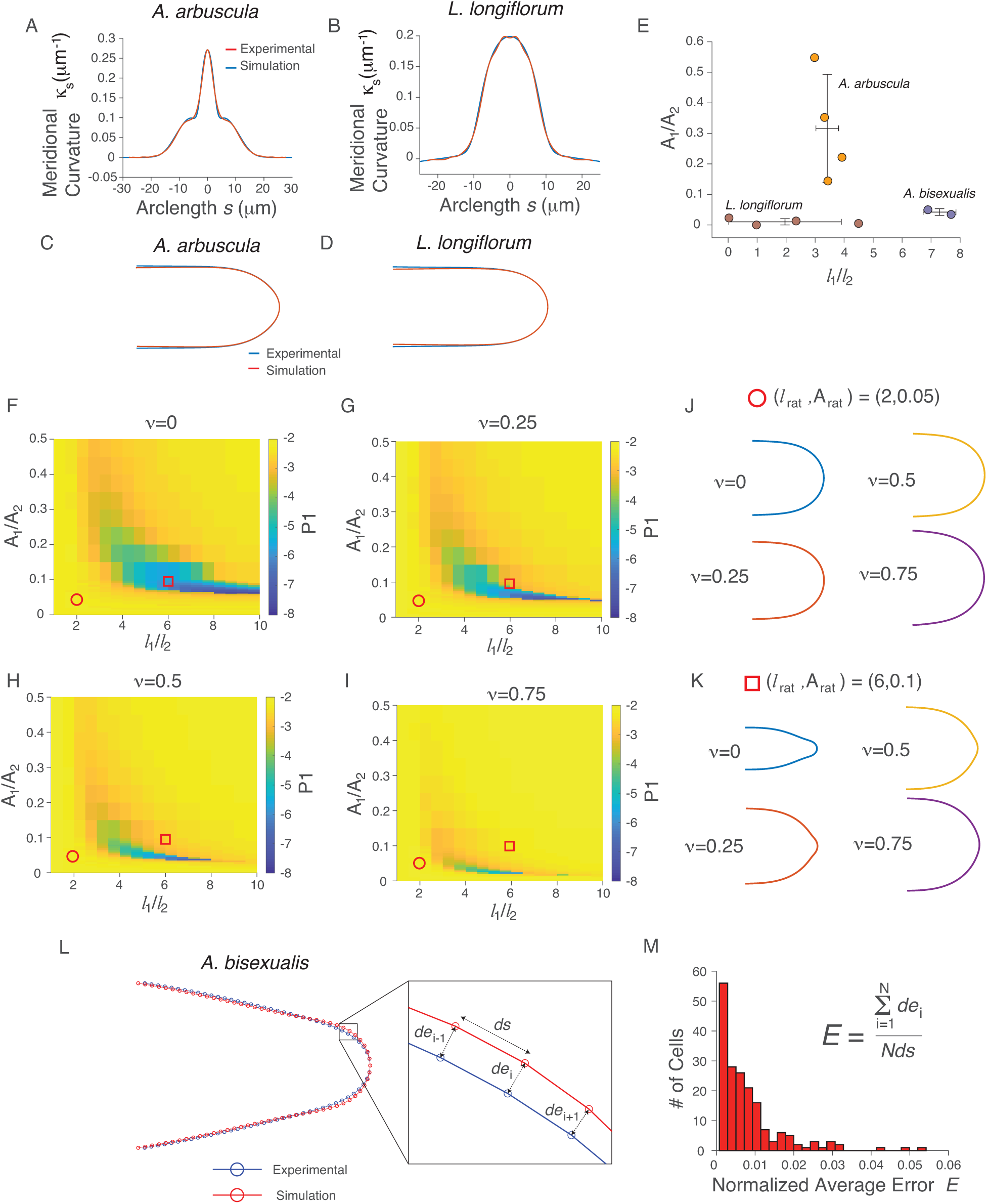
The Gaussian mixture surface extensibility profile predicts apical geometry. (A,B) Mean meridional curvature versus arclength for cells from *A. arbuscula* and *L. longiflorum* cells (blue line, same data shown in Fig. 2D of main text), and the curvatures of cells simulated using a transverse isotropy model and a Gaussian mixture surface extensibility profile with *l*_rat_ and *A*_rat_ that yielded the best fit of the experimental surface extensibility profile (red line, Fig. 3A-C of the main text). (C,D) Time-averaged apical geometries for *A. arbuscula* and *L. longiflorum* cells (blue line), and the apical geometries of cells simulated using a transverse isotropy model with a Gaussian mixture surface extensibility profiles with *l*_rat_ and *A*_rat_ that yielded the best fits of the experimental extensibility profile (red lines). (E) Values of (*l*_rat_,*A*_rat_) that yielded the best fit of the experimental surface extensibility profile versus species. Error bars indicate ± 1 s.d. (F-I) P1 of simulated cells using the transverse isotropy model and a Gaussian mixture surface extensibility profile, as a function of *l*_rat_ and *A*_rat_, for four different values of *v*. (J) Apical geometries of simulated cells for four different values of *v* at at point in morphospace where *v* does not modulate geometry, (*l*_rat_, *A*_rat_) = (2, 0.05), indicated by the red circle in (F-I). (K) Apical geometries of simulated cells for four different values of *v* at at point in morphospace where *v* greatly modulates geometry, (*l*_rat_,*A*_rat_) = (6, 0.1), indicated by the red square in (F-I). (G) Time-averaged apical geometry for an *A. bisexualis* cell (blue line) and the apical geometry of a cell simulated using the transverse isotropy model and a Gaussian mixture surface extensibility profile with *l*_rat_ and *A*_rat_ that yielded the best fit of the experimental extensibility profile (red line, Fig. 3A). (inset) The error associated with the simulation is calculated for each point used to discretize the outlines. (H) The normalized average error, *E*, for the cell is equal to the average error for each point used to discretize the outlines, divided by the interpoint arclength, *ds*.

